# Effects of irrigation scheduling and different irrigation methods on onion and water productivity in Tigray, northern Ethiopia

**DOI:** 10.1101/790105

**Authors:** Gebremedhin Gebremeskel Haile, T.G. Gebremicael, Mulubrehan Kifle, Teferi Gebremedhin

**Author notes:** Corresponding author, (GGH).

## Abstract

Efficient irrigation water use can meet its objective if irrigated agriculture is managed properly in regions where water is limited. A two-year (2016-2017) field experiments were conducted in the semiarid areas of Tigray in northern Ethiopia to evaluate irrigation scheduling with different irrigation methods. The experiments were carried out to identify their contribution for enhancing onion and water productivity in water-stressed irrigation schemes of Korir and Hatset sites. Six factorial treatments comprising of three levels of irrigation methods (furrow, basin and flood) and two levels of irrigation scheduling (fixed interval and farmer’s practices) were evaluated with three replications. The agronomic and irrigation parameters were subjected to separate level-wise comparison followed by the factorial interaction effects. The results showed that the fixed irrigation interval, basin irrigation method and their factorial combinations showed better performances and produced a higher yield and water productivity. On average, 263.85 q/ha and 281 q/ha of onion were obtained under the basin irrigation method and basin irrigation with fixed irrigation interval (T2) at both sites and seasons. For the T2, 6.27 and 6.06 kg/m^3^ of water productivity (WP) and 4.39 and 4.24 kg/m^3^ of irrigation water productivity (IWP) were obtained at Korir and Hatset sites, respectively showing higher results as compared to other treatment combinations. Moreover, the basin irrigation method produces higher marketable onion bulbs that are essential for onion producers to earn maximum profit from selling and enhance their livelihoods. Treatments comprising basin method and fixed interval showed better onion and water productivity in the water-limited irrigation schemes of Tigray. Hence, focusing on enhanced irrigation scheduling techniques and irrigation methods are important for effective agricultural water management. Farmers, irrigation experts, water resources managers and decision-makers are suggested to use these techniques to save the limited water resources and increase agricultural productivity.

## Introduction

Agriculture has been the basis of the Ethiopian economy for centuries [1,2]. Agricultural productivity was growing continuously in the last two decades, even though the population was also increasing over time [3–6]. Water resources have played a vital role for increased agricultural production especially in the semiarid areas of northern Ethiopia [7–10]. However, water is being threatened by various environmental and socioeconomic variabilities [11–14]. Water scarcity is likely to increase significantly as the requirement for food production and industrial use is at an increasing rate [15]. The attention given for improving agricultural water management is also non-sufficient [15]. This clearly suggests that enhancing water productivity is important in poverty reduction and future economic development endeavours in the semiarid areas of the country. Efficient water utilization based on the crop water requirement accompanied with optimum crop productivity is also needed to sustain economic developments.

In water-stressed regions (e.g., irrigation areas in northern Ethiopia), efficient irrigation water use can meet its objective if the agricultural field is managed properly. However, improper on-farm irrigation management practices could lead to poor water distribution; non-uniform crop growth; waterlogging and aggravates salinity build-ups [10,14–18]. Literature show that farmers who practice poor water management techniques obtain low- and poor-quality irrigation products such as onion yields with a maximum loss of water and labour forces [15,16,19]. To this effect, efficient irrigation water application techniques substantiated with appropriate field management strategies are required to improve water productivity in water scares areas. There are various widely applied irrigation methods (e.g., basin, furrow and flooding) and irrigation scheduling techniques in the region [10,16]. However, the knowledge on the linkage of these irrigation water application methods and irrigation scheduling techniques is lacking. As a result, understanding the combined effects of irrigation methods and irrigation interval can be used to enhance effective agricultural water management in irrigated agriculture [16,20,21]. Besides, evaluating these irrigation techniques is vital to put recommendations for irrigation users for optimum crop production.

Onion (*Allium cepa* L.) crop is one of the most dominantly consumed cash crops in Ethiopia [16,22–24]. It is a key component in the Ethiopian daily foodstuff as it improves the taste and scent of the food [22]. In addition, onion is consumed more than any other vegetable crops in Ethiopia and 95% of its production is produced by smallholder farmers [25]. Despite its importance, onion productivity in the region remains less than 100 q/ha which is very low as compared to the above 197 q/ha world’s average production [26]. This has been drawing the attention of researchers for increasing the productivity of onion crop in the region via introducing efficient irrigation water management strategies. Several studies [27–30] showed that among the different irrigation methods, drip and sprinkler irrigation methods increases the yield of onion and water productivity. However, due to lack of technical knowledge, high labour demand, and poor extension services, these irrigation methods are not well adapted and farmers are distrustful to their acceptance [14].

In northern Ethiopia (Tigray region), most farmers use conventional irrigation practices instead of the drip and sprinkler irrigation systems to irrigate their crops [16,18,31]. As a result, these mechanized irrigation practices (drip irrigation and sprinkler) are limited in the region [10,14,17]. Some attempts have been carried out to introduce and popularized drip irrigation as efficient water-saving technology in the region. However, it was not successful to be adopted by the farmers due to various reasons as discussed in Gebremeskel et al. [14] and Haregeweyn et al. [32]. Traditional irrigation practices are known to be less efficient particularly when there is a scarcity of irrigation water [33]. Surface irrigation methods are commonly practised by the farmers in most irrigation schemes of the region. However, a uniform production system is hardly found because of the knowledge gaps between producers and extension experts [15]. These differences are clearly seen in the irrigation methods, the frequency of irrigation and plant spacing of onion cultivation processes under irrigation condition. The extension service recommends furrow irrigation method rather than others, whereas farmers prefer to use a basin to produce onion. Although various studies [16,17,24,25,33] have conducted in the region on onion production and productivity, there is no clear understanding on which method of irrigation and watering frequency gives better quantity and quality of onion. Therefore, this study aimed at investigating the effect of different irrigation methods and watering frequency which could maximize the yield and productivity of onion. Furthermore, evaluating different irrigation methods and irrigation interval along with their interaction are important for the current irrigation development endeavours to maximize the productivity of water and increase the productivity of onion in the region.

The paper is structured as follows. Following this introduction, study area description, experimental setup and data analysis including the agronomic and irrigation scheduling strategies are presented in the materials and methods section. In the results and discussion section, graphical and tabular results and corresponding detailed discussions are presented. Finally, concluding remarks are given under the conclusion section.

## Materials and methods

### Study area

This study was conducted at Korir and Hatset irrigation sites, which are found in Kilte Awlaelo and Hawzen districts of the Eastern Tigray region in Northern Ethiopia (Fig 1). The Korir experimental site is located at 13.64°N and 39.59°E and 2059 m a.s.l while Hatset is located between 14.06°N and 39.49°E at 1935 m a.s.l. The sources of water for irrigation in Korir and Hatset schemes is from earthen dams constructed in 1997 and 2015, respectively [14,31]. Subsistent small-scale farmers have predominantly use irrigation schemes to produce crops during the long dry season (November-June). Agricultural land is the dominant land use/cover, followed by bushland and rural settlements in both irrigation schemes [34,35]. Sandy clay loam and sandy loam are the dominant soil textural types in Korir and Hatset irrigation schemes, respectively. In the areas where both irrigation schemes are located, the rainfall is uni-modal characterized by high variability [19,36]. Such variabilities are mainly associated with the seasonal migration of the inter-tropical convergence zone (ITCZ) and its complex topography [37–40]. The annual rainfall and evapotranspiration of both sites range from 400-550 mm and 1400-1600 mm, respectively [10,19].

**Fig 1.**
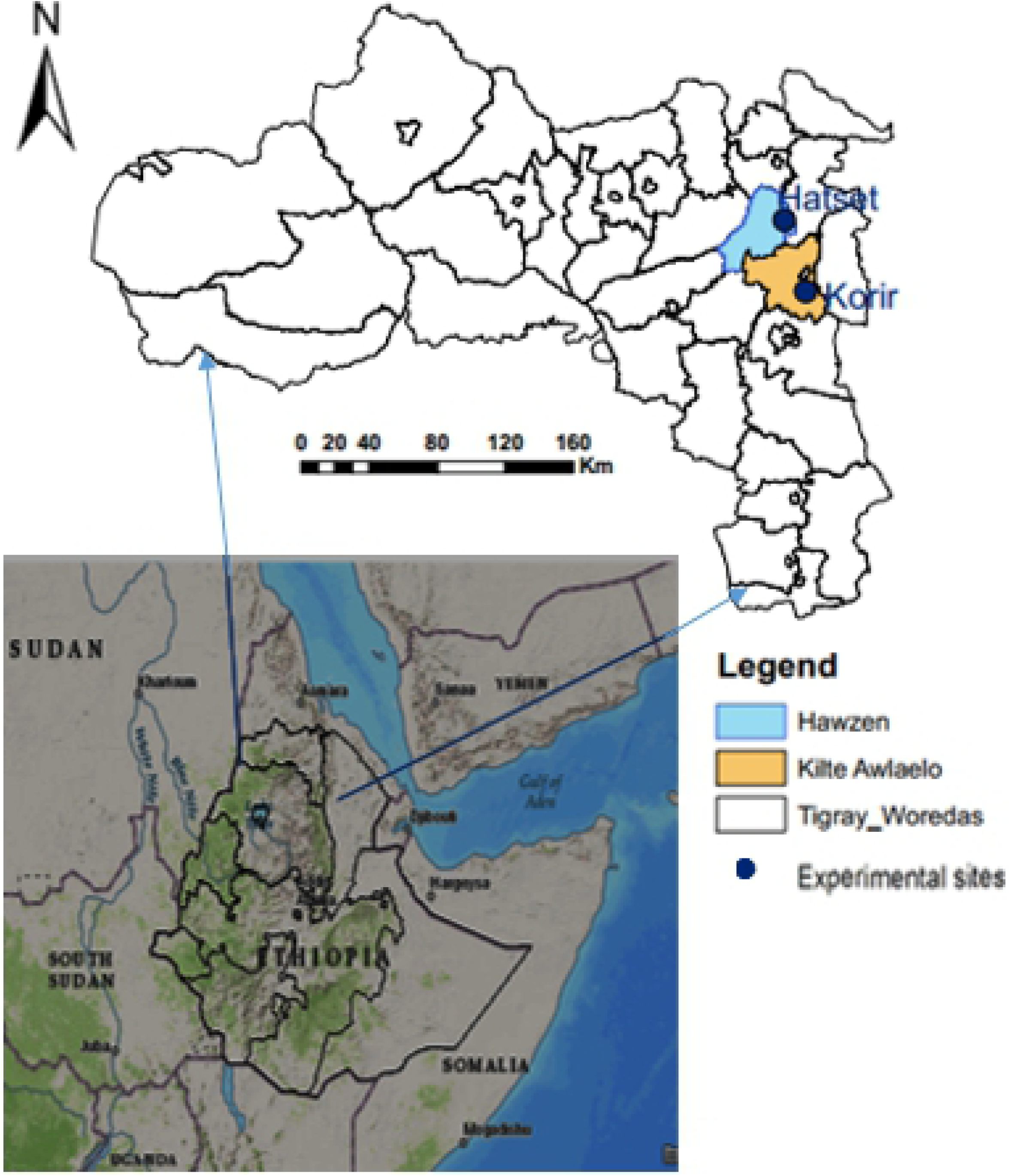
Map of the study area.

### Treatment setup and experimental design

The experiment was laid in a randomized complete block design replicated three times at two experimental sites (Korir and Hatset). The experiments were conducted for two successive years (2016 and 2017). The experiments consist of two factors, namely irrigation scheduling practices (fixed interval and farmers’ practice) and irrigation methods (furrow, basin and flood). The fixed irrigation intervals were computed based on the crop water requirement whilst the farmers’ practice was a locally available irrigation interval practised by farmers. Six treatment combinations were randomly distributed in each experimental site (Table 1). Each treatment has a plot size of 4 m × 4 m and the layout of the experiment is presented in Fig 2. For the furrow irrigation method, a 40 cm spacing was applied. For the remaining, the plant spacing was 10 cm with 20 cm apart in a double row. The spacing between blocks and treatments was maintained to be 1.25 m and 0.75 m, respectively. Therefore, the total area needed for a single experiment was 420 m^2^ (28m × 15m). Such size of the treatment plots was determined considering the farmer’s landholding size as most farmer’s in the region owns very small plots. To this effect, most experimental irrigation studies have been applied an experimental plot size of less than 4 × 4m [15,30]. Using these specifications, the experiments were undertaken for two irrigation seasons at two sites. The same experimental plots were applied in both seasons with similar treatment setups.

**Fig 2.** Layout of the experiment (note: p stands for a plot)

**Table 1.** Treatments setup.

| **Irrigation interval** | **Irrigation methods** | **Treatment combinations** | **Representations** |
| --- | --- | --- | --- |
| **Fixed irrigation (Fi)** | Flood irrigation (F) | FiF | T6 |
|  | Furrow irrigation (Fr) | FiFr | T4 |
|  | Basin irrigation (B) | FiB | T2 |
| **Farmers' practice (Fp)** | Flood irrigation (F) | FpF | T5 |
|  | Furrow irrigation (Fr) | FpFr | T3 |
|  | Basin irrigation (B) | FpB | T1 |

### Data collection and analysis

Climate and soil data are required to compute crop water requirement and irrigation scheduling using FAO CROPWat software [41]. The long-term average climatic information of the irrigation sites for the experimental months was collected from nearby meteorological stations. A composite of soil samples from 0-20 cm depth, where the maximum rooting depth of onion reaches was collected to characterize the physical and chemical properties of the soil. The samples were collected using soil auger from both sites. Soil parameters of the collected samples were analysed at Mekelle Soil Research Centre’s laboratory following the standard procedures given by Sahlemedhin and Taye [42].

### Crop water requirement and irrigation scheduling

Crop water requirement of onion for the growing seasons at both sites was computed using reference evapotranspiration estimated from climatic data and crop coefficient of onion. As there is no site-specific crop coefficient in the region, the different values for the initial, middle and late growth stages of onion were taken from Allen et al., [41] and hence crop water requirement (CWR) was calculated using Equation 1. Considering the experimental sites’ irrigation experience and existing research outputs from the irrigation sites [16], 70% of field efficiency was applied for computing the gross CWR in both seasons and sites. Irrigation scheduling of onion was determined by considering soil type of each site and fixed interval and variable depth (refill to field capacity). Two levels of watering intervals, i.e. fixed interval and farmers practice were employed to compute irrigation scheduling of onion. In the fixed irrigation interval, irrigation water was applied based on the crop water requirement of the crop whilst in farmers’ practice irrigation interval, locally available fixed intervals were applied in both sites.

For the fixed irrigation interval, the CWR of onion at each irrigation site were estimated following the procedures given in Equations 1-3. The CWR and irrigation scheduling was computed using the formula given in Equation 1. Based on the soil, climatic and local irrigation experiences in the experimental sites, seven days irrigation interval at Korir and six days at Hatset was considered to schedule the application of water. The amount of water applied to each plot was measured at every irrigation time using Parshall flume [43,44].

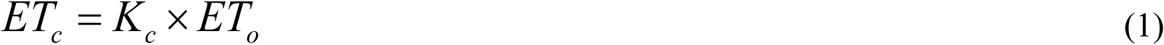

Where; ET_c_ the crop water requirement, K_c_ is the crop coefficient, and ET_o_ is the reference evapotranspiration.

As the amount of rainfall received during the experimental months was negligible, the net irrigation requirement was the same as ET_c_. The irrigation water productivity (IWP) and the total crop water productivity (WP) was calculated as given in Equations 2 and 3, respectively [42,45].

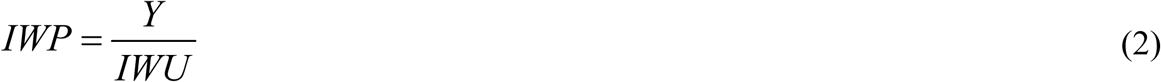

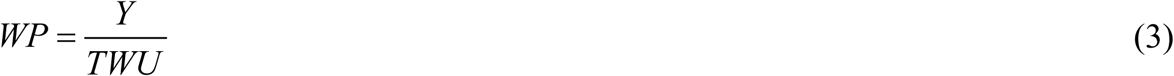

Where; IWP is in kg/m^3^, IWU is irrigation water use (mm), WP is water productivity (Kg/m^3^), TWU is the total water use, and Y is the bulb yield (kg/ha)

In the farmers’ irrigation interval, measurements for the irrigation applications from farmers’ practice was performed at the time of irrigation using Parshall flume [43,44]. Required data were taken at each irrigation time from both irrigation sites to estimate the irrigation water and intervals applied to the onion. The average of the measurements and the total number of irrigation intervals were considered to estimate the irrigation water productivity of onion in both farmer’s practices and designed experimental plots. To irrigate the onion crop farmers regularly use a constant interval to irrigate crops based on their specific observation on the performance of onion leaves. Traditionally, farmers use to observe the leaves of onion crop in order to decide the next irrigation period. That is, when the colour of the leaves turns from green to yellow, farmers decide to irrigate their crops. This has been a common traditional irrigation practice/system in the region that we are interested to take into account in this experiment.

### Agronomic management

In this study, a widely used Bombay red onion variety (*Allium cepa* L.) was used as an indicator crop. After we have raised this variety at three beds (1m × 4m) with the seed rate of 4kg/ha, the seedlings were transplanted to the experimental sites 50 days after growing in the seedbeds. Di-Ammonium Phosphate (DAP, 200 kg/ha) comprising 18N-46P fertilizers and urea (100 Kg/ha) was applied to the treatment plots following the recommendations in the irrigation schemes. DAP was applied at the time of planting whereas urea was applied half at the time of planting and the remaining was top-dressed three weeks after planting. Each plot received the same fertilizer dose and field management in both years and sites. Moreover, standard methods of fertilizer application systems and weed management strategies were used in the experiments.

### Statistical analysis

Collected data including agronomic and irrigation water application parameters of onion were subjected to statistical analysis using GenStat statistical software [46]. Treatments mean differences were determined using least significant difference (LSD) comparison method at 95% confidence level [47]. The coefficient of variation (CV) was also calculated to identify the relative variability among the treatments. Moreover, the irrigation intervals were analysed to determine if there are variations between the fixed irrigation interval and farmers’ irrigation interval and were subjected for analysis using error bars. Error bars are graphical representations of the variability of data which are commonly used to show the uncertainties in a given experimental measurement [48]. An independent analysis was first undertaken for irrigation intervals (fixed interval and farmers practice) and irrigation methods (basin, furrow and flood irrigation) separately and then combined/factorial analysis were followed for the treatment combinations for each site and each irrigation season. Furthermore, a combined analysis was also applied by taking the average of the two-year data for each site to compare and further investigate possible statistical variations among the treatments considered.

## Results and discussion

### Soil and climate characteristics

#### Soil physicochemical properties of the irrigation schemes

Summary of analysed soil physical and chemical properties of the irrigation sites is given in Table 2. Accordingly, sandy clay loam and sandy loam soil texture were found Korir and Hatset sites, respectively. The experimental sites have field capacity (FC) of 25.06% and 23.2% and permanent wilting point (PWP) of 11.16% and 12.6% for Korir and Hatset, respectively. The average total available water (TAW) by volume percentage is also estimated as 139 and 106% for Korir and Hatset, respectively.

**Table 2.** Characteristics of the experimental soil (0-20 cm, composite soil).

| Soil characteristic parameters | Experimental sites |  |
| --- | --- | --- |
|  | Korir | Hatset |
| Sand (%) | 51 | 59 |
| Silt (%) | 27 | 23 |
| Clay (%) | 22 | 18 |
| Texture | Sandy clay loam | Sandy loam |
| Soil pH | 7.24 | 6.89 |
| Electrical conductivity (dS/m) | 0.77 | 0.82 |
| Organic matter (%) | 0.9 | 0.95 |
| Bulk density (g/cm<sup>3</sup>) | 1.42 | 1.48 |
| Field capacity vol. (%) | 25.06 | 23.2 |
| Permanent wilting point vol. (%) | 11.16 | 12.6 |
| TAW (mm/m) | 139.10 | 106.0 |

#### Climate characteristics of the irrigation schemes

The climatic data including, precipitation, maximum and minimum temperature, humidity, sunshine hours and solar radiation were obtained from nearby stations, Hawzen and Wukro for Hatset and Korir, respectively. Table 3 presents the summary of climate information for both irrigation sites. Each of the basic climatic characteristics of the schemes are given as long-term monthly averages. The amount of rainfall received during the growing seasons was negligible in both sites (Tables 3-5).

**Table 3.** Long-term monthly average climatic characteristics of the irrigation sites.

| Experimental sites | Crop growth months | Temperature (°C) |  | Humidity (hPa) | Wind speed (Km h<sup>-1</sup>) | Sunshine hours (%) |
| --- | --- | --- | --- | --- | --- | --- |
|  |  | Minimum | Maximum |  |  |  |
| **Korir** | **February** | 7.5 | 25.4 | 9.05 | 5.6 | 81.8 |
|  | **March** | 10.2 | 26.9 | 9.82 | 6.5 | 78.7 |
|  | **April** | 12.3 | 26.4 | 10.95 | 6.5 | 76.2 |
|  | **May** | 12.5 | 26.3 | 10.91 | 7.5 | 80.5 |
| **Hatset** | **February** | 6.5 | 24.4 | 10.11 | 5.76 | 77.5 |
|  | **March** | 7.3 | 26.5 | 10.20 | 6.12 | 81.4 |
|  | **April** | 10.5 | 27.8 | 11.23 | 6.15 | 80.9 |
|  | **May** | 9.5 | 25.7 | 11.95 | 7.5 | 87.2 |

### Crop water requirement and irrigation scheduling of onion

The amount of irrigation water requirements, irrigation events and irrigation scheduling at each site has been summarized in Table 4 (for Korir) and Table 5 (for Hatset). The growing period and gross irrigation water requirement of onion are estimated to be 98 days and 607.9 mm at Korir and 95 days and 600.9 mm at Hatset, respectively. Similarly, for the farmers’ practice, irrigation water requirement was obtained simply by measuring the amount of water they applied at each time of watering. As a result, the gross irrigation water applied by the farmers in both experimental sites were estimated to be 780 and 856 mm at Korir and Hatset, respectively. The irrigation interval for the farmer’s practices treatments was 5 and 6 days, in Korir and Hatset, respectively. The irrigation intervals used by the farmers in Korir site is much closer than the farmers from Hatset as compared to the CROPWat results from each site (Table 4 and 5). This indicates that farmers in Korir irrigation scheme have developed better irrigation experiences linked to the age of earthen dam as they have used it for the last two decades. Farmers in Haste, however, use irrigation more frequently as compared to the interval estimated by the CROPWat and the lower relative experiences they have to irrigation water use. These suggest that farmers are tending to use excess irrigation water to irrigate their onion crop in this scheme.

**Table 4.** Crop water requirement and irrigation scheduling of onion at Korir irrigation scheme for the year 2017.

| **Date** | **Interval**<br>**(days)** | **ET<sub>o</sub>**<br>**(mm)** | **K<sub>c</sub>** | **ET<sub>c</sub>**<br>**(mm)** | **P<sub>eff.</sub>**<br>**(mm)** | **IR<sub>n</sub>**<br>**(mm)** | **IR<sub>g</sub>**<br>**(mm)** |
| --- | --- | --- | --- | --- | --- | --- | --- |
| **14-Feb** | 7 | 27.72 | 0.70 | 19.4 | 0.00 | 19.4 | 27.7 |
| **21-Feb** | 7 | 27.72 | 0.70 | 19.4 | 0.00 | 19.4 | 27.7 |
| **28-Feb** | 7 | 27.72 | 0.70 | 19.4 | 0.00 | 19.4 | 27.7 |
| **7-Mar** | 7 | 32.48 | 0.78 | 25.3 | 0.00 | 25.3 | 36.2 |
| **14-Mar** | 7 | 32.48 | 0.84 | 27.3 | 0.00 | 27.3 | 39.0 |
| **15-Mar** | 7 | 32.48 | 0.89 | 28.9 | 0.00 | 28.9 | 41.3 |
| **21-Mar** | 7 | 32.48 | 1.01 | 32.8 | 0.00 | 32.8 | 46.9 |
| **28-Mar** | 7 | 32.48 | 1.01 | 32.8 | 0.00 | 32.8 | 46.9 |
| **4-Apr** | 7 | 35.70 | 1.05 | 37.5 | 0.00 | 37.5 | 53.6 |
| **11-Apr** | 7 | 35.70 | 1.05 | 37.5 | 0.00 | 37.5 | 53.6 |
| **18-Apr** | 7 | 35.70 | 1.05 | 37.5 | 0.00 | 37.5 | 53.6 |
| **25-Apr** | 7 | 35.70 | 1.05 | 37.5 | 0.00 | 37.5 | 53.6 |
| **2-May** | 7 | 35.84 | 1.00 | 35.8 | 0.00 | 35.8 | 51.2 |
| **9-May** | 7 | 35.84 | 0.96 | 34.4 | 0.00 | 34.4 | 49.2 |
| **Total** | 98 | 460.04 |  | 425.53 |  | 425.53 | 607.90 |

ET<sub>o</sub>= Reference evapotranspiration, K<sub>c</sub>= Crop coefficient, ET<sub>c</sub>= Crop evapotranspiration, P<sub>eff</sub>=Effective rainfall, IR<sub>n</sub>= Net irrigation requirement, IR<sub>g</sub>= Gross irrigation requirement.

**Table 5.**
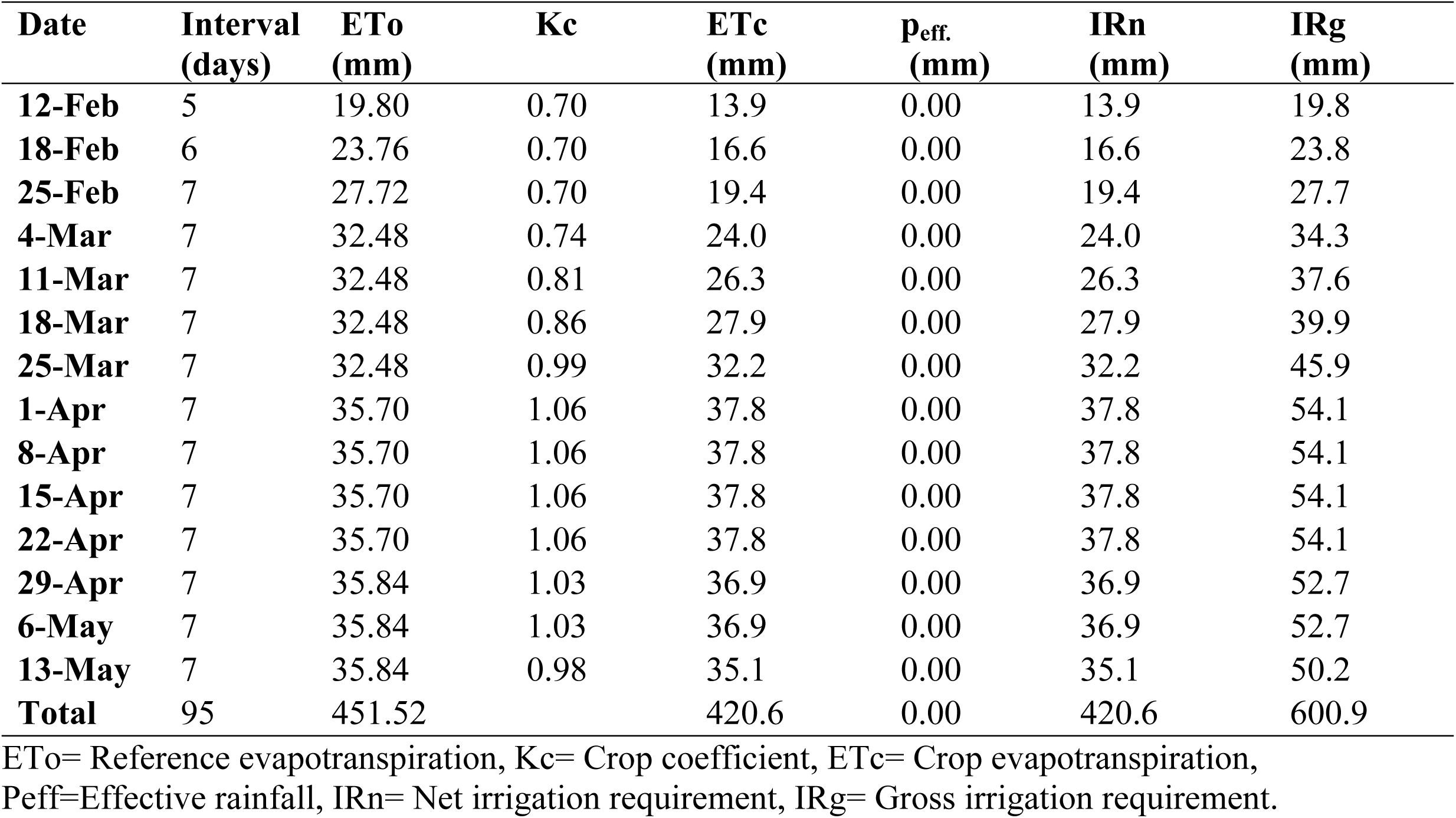
Crop water requirement and irrigation scheduling of onion at Hatset irrigation scheme.

### Effects irrigation intervals and different irrigation methods on onion yield

#### Effects of irrigation intervals on onion yield

Fig 3 presents the average yield parameter obtained from both growing seasons in both experimental sites. The error bars shown in each graph indicates the variabilities and uncertainties of the data which clearly shows how spread the data are around the mean value. As shown in Fig 3, there is no overlap between the standard deviation errors bars which imply that the difference is not statistically significant. This indicates that irrigation intervals showed non-significant effects on the total yield of onion in both experimental sites (Fig 3). Similarly, the marketable yield was not significantly influenced by the effects of the irrigation intervals.

**Fig 3.**
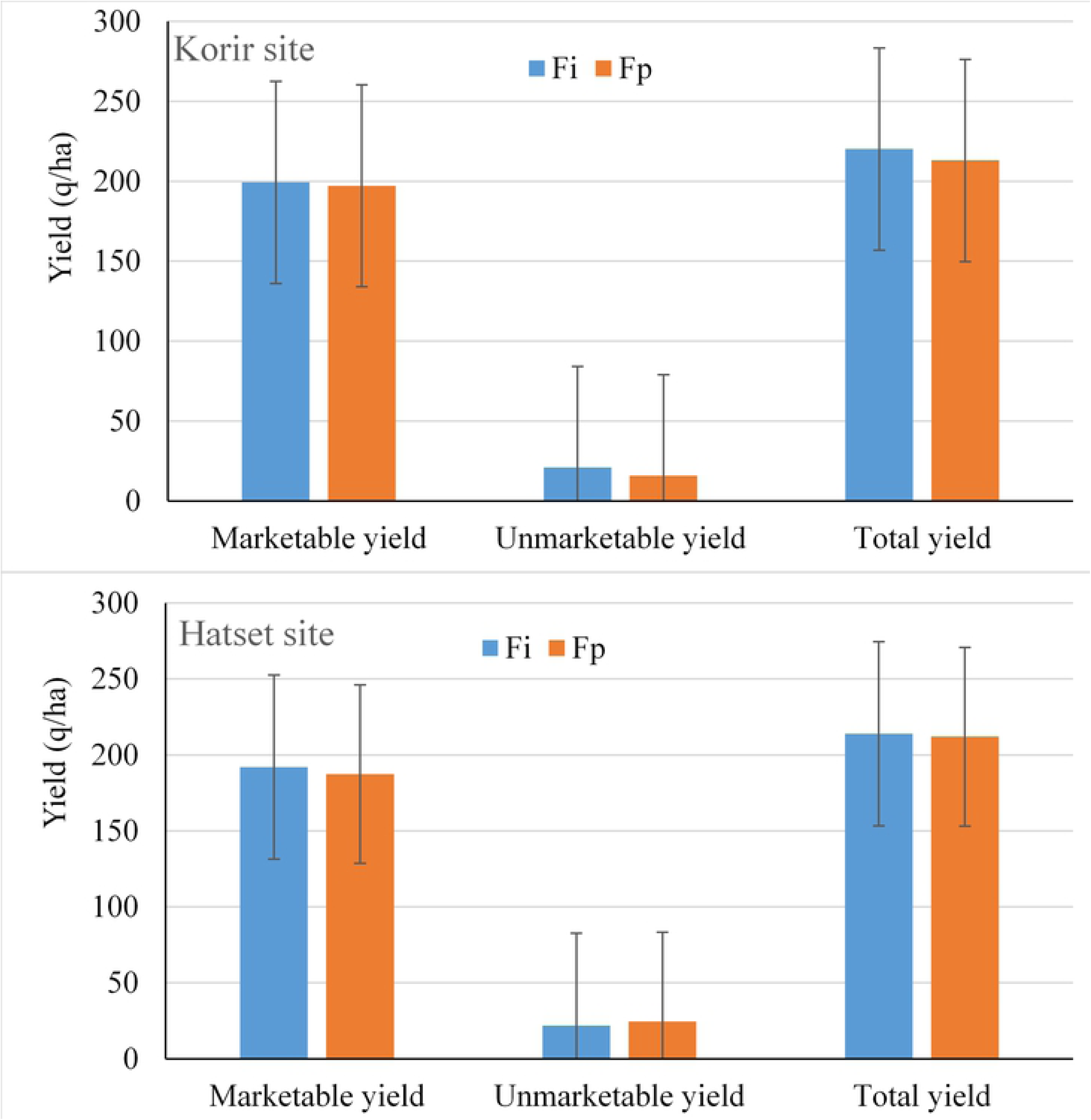
Effects of irrigation interval on onion yield in 2016 and 201.

It is worthwhile to explain that the average yield of onion from both sites is comparable. However, despite statistically non-significant, a relatively better yield was obtained from the fixed interval treatments from both sits. The marketable yield is found to be 199.22 and 191.94 q/ha under Fi and 197.14 and 187.36 q/ha under Fp for Korir and Hatset sites, respectively. Similarly, the bulb yield is found to be 220.13 and 213.89 q/ha for Fi, and 212.92 and 211.94 q/ha for Fp at Korir and Hatset sites, respectively. Higher yields of both marketable and bulb yield were observed from Fi irrigation interval as compared to the Fp.

#### Effects different irrigation methods on onion yield

The effects of irrigation methods (basin, furrow, and flooding) on the yield of onion in the two experimental sites (Korir and Hatset) are summarized in Table 6. The results showed that the total bulb yield and marketable yields have shown a significant difference with the irrigation methods. The basin irrigation method was significantly higher than the furrow and flood irrigation methods at a 5% significance level in all conditions. The Korir irrigation scheme had the highest onion marketable yield (245.1 q/ha) under basin irrigation and the least yield was obtained from the flood irrigation treatment (162.3 q/ha) in 2016. A similar pattern was observed in the 2017 growing season. Next to basin irrigation method, the higher yield was obtained from flooding, and furrow irrigation methods, respectively (Table 6). However, the differences between flooding and furrow irrigation methods were not statistically significant. Table 6 also presents the effect of the irrigation methods on the average of the two-year results from both growing seasons on marketable and bulb onion yields. The basin irrigation method continued to show a higher marketable and bulb yield at both sites. Both independent (year wise) and average yield analysis from both growing seasons showed that the basin irrigation method is strong enough in producing a higher bulb yield compared to the flooding and furrow methods.

**Table 6.**
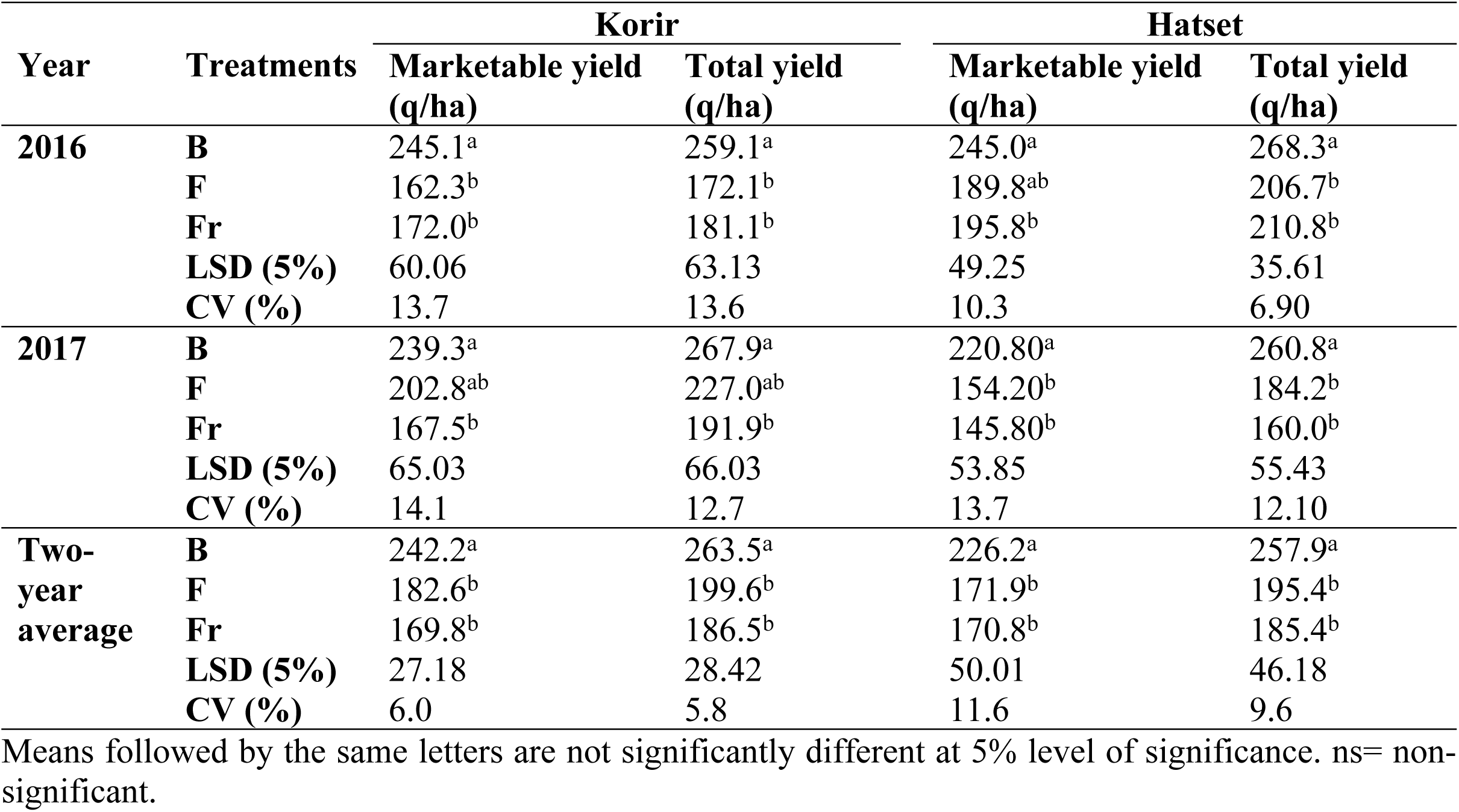
Effects of irrigation methods on the marketable and bulb yield of onion.

#### Treatment interaction effects on onion yield

Marketable and total yield were significantly influenced by the interaction effect of irrigation methods and irrigation interval in both irrigation schemes (Table 7). For example, during the 2016 season, the highest and lowest marketable and total yields of onion were obtained from treatment combinations of T1 and T6 at Korir irrigation scheme. Statistically, significant differences among the treatment combinations were found when the yield is averaged (Table 7). In contrast, a higher and lower marketable yield were obtained under T2 and T4 combinations in Korir irrigation scheme. This is different in Hatset irrigation scheme where the maximum and minimum yield were obtained from treatments T1 and T3 (Table 7).

**Table 7.** Factorial effects of irrigation methods and irrigation interval on onion yield at Korir and Hatset sites.

| Year | Treatment combinations | Korir |  | Hatset |  |
| --- | --- | --- | --- | --- | --- |
|  |  | Marketable yield (q/ha) | Total yield (q/ha) | Marketable yield (q/ha) | Total yield(q/ha) |
| 2016 | T1 | 254.2<sup>a</sup> | 265.8<sup>a</sup> | 240.0<sup>a</sup> | 256.7<sup>a</sup> |
|  | T2 | 235.9<sup>ab</sup> | 252.4<sup>a</sup> | 223.3<sup>ab</sup> | 253.3<sup>ab</sup> |
|  | T3 | 177.5<sup>bc</sup> | 184.8<sup>b</sup> | 202.5<sup>ab</sup> | 215.0<sup>bc</sup> |
|  | T4 | 166.6<sup>c</sup> | 177.4<sup>b</sup> | 189.2<sup>b</sup> | 206.7<sup>c</sup> |
|  | T5 | 176.6<sup>bc</sup> | 185.9<sup>b</sup> | 181.7<sup>b</sup> | 201.7<sup>c</sup> |
|  | T6 | 148.1<sup>c</sup> | 158.3<sup>b</sup> | 197.5<sup>ab</sup> | 211.7<sup>c</sup> |
|  | LSD (5%) | 63.13 | 63.99 | 47.45 | 39.26 |
|  | CV (%) | 18.0 | 17.2 | 12.7 | 9.6 |
| 2017 | T1 | 227.6<sup>ab</sup> | 254.4<sup>ab</sup> | 216.7<sup>a</sup> | 265.0<sup>a</sup> |
|  | T2 | 251.0<sup>a</sup> | 281.4<sup>a</sup> | 225.0<sup>a</sup> | 256.7<sup>a</sup> |
|  | T3 | 164.1<sup>b</sup> | 184.8<sup>b</sup> | 141.7<sup>b</sup> | 155.0<sup>b</sup> |
|  | T4 | 170.9<sup>ab</sup> | 198.9<sup>b</sup> | 150.0<sup>b</sup> | 165.0<sup>b</sup> |
|  | T5 | 182.8<sup>ab</sup> | 201.7<sup>b</sup> | 141.7<sup>b</sup> | 178.3<sup>b</sup> |
|  | T6 | 222.8<sup>ab</sup> | 252.2<sup>ab</sup> | 166.7<sup>ab</sup> | 190.0<sup>b</sup> |
|  | LSD (5%) | 83.97 | 78.06 | 62.5 | 59.44 |
|  | CV (%) | 22.7 | 18.7 | 19.8 | 16.2 |
| Two-year average | T1 | 240.9<sup>a</sup> | 260.1<sup>a</sup> | 228.3<sup>a</sup> | 260.8<sup>a</sup> |
|  | T2 | 243.5<sup>a</sup> | 266.9<sup>a</sup> | 224.2<sup>ab</sup> | 255.0<sup>a</sup> |
|  | T3 | 170.8<sup>b</sup> | 184.8<sup>b</sup> | 172.1<sup>c</sup> | 185.0<sup>b</sup> |
|  | T4 | 168.7<sup>b</sup> | 188.2<sup>b</sup> | 169.6<sup>c</sup> | 185.8<sup>b</sup> |
|  | T5 | 179.7<sup>b</sup> | 193.8<sup>b</sup> | 161.7<sup>c</sup> | 190.0<sup>b</sup> |
|  | T6 | 185.5<sup>b</sup> | 205.3<sup>b</sup> | 182.1<sup>bc</sup> | 200.8<sup>b</sup> |
|  | LSD (5%) | 51.33 | 48.03 | 44.14 | 39.59 |
|  | CV (%) | 14.2 | 12.2 | 12.8 | 10.2 |

Means followed by the same letters are not significantly different at 5% level of significance. ns= non-significant

Comparing to flooding and furrow irrigation methods, treatment combinations having basin irrigation methods showed higher yields. T1 and T2 revealed a higher bulb and marketable yields in both experimental sites when the irrigation methods are associated with the irrigation intervals.

### Effects of irrigation intervals and irrigation methods on yield components of onion

The effects of irrigation interval and irrigation methods on yield components of onion were evaluated separately and in combination with the individual factors. A separate result of the individual levels and their interactions are given in Fig 4, Table 8 and 9.

**Fig 4.**
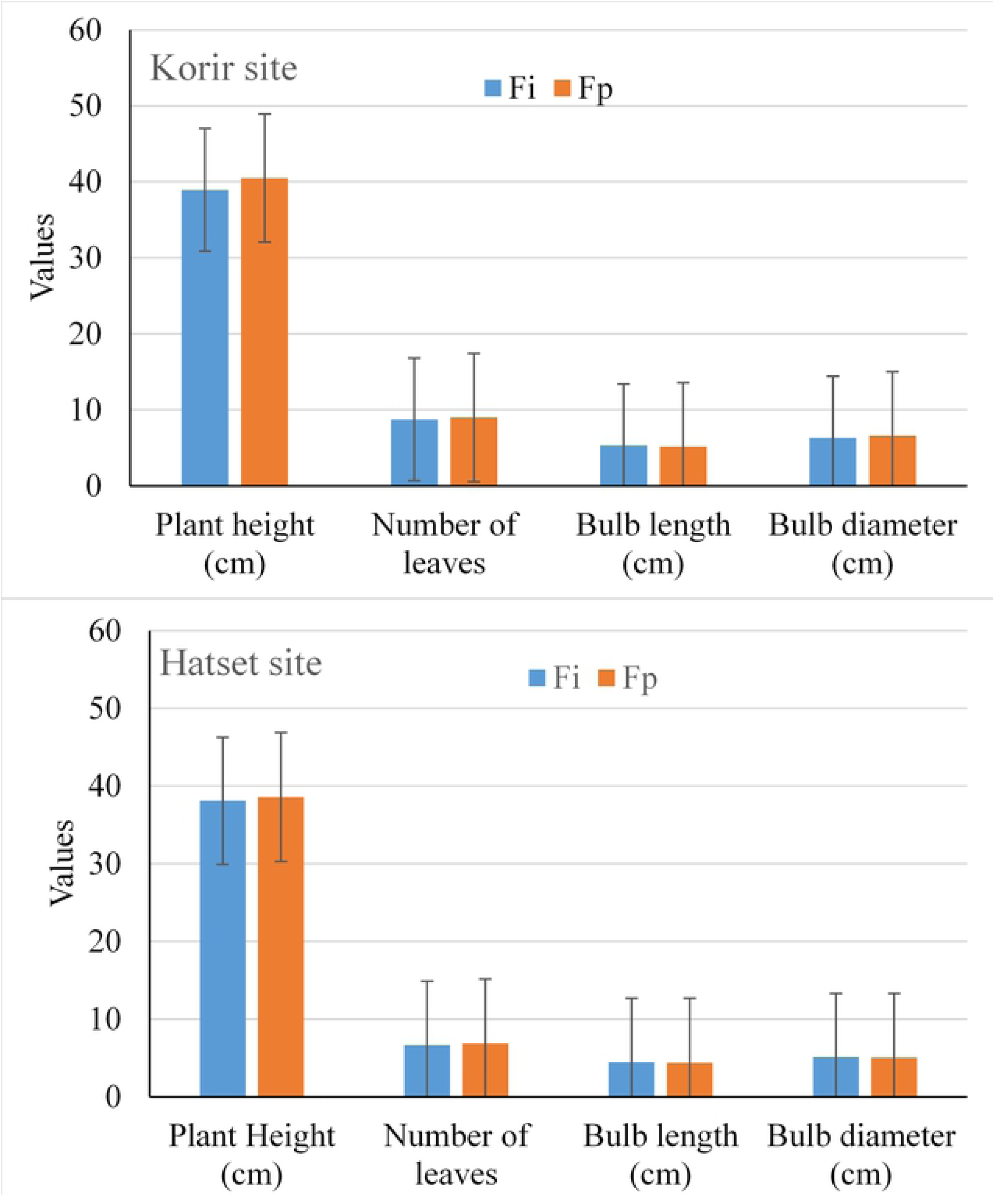
Effects of irrigation interval on onion yield components in.

**Table 8:**
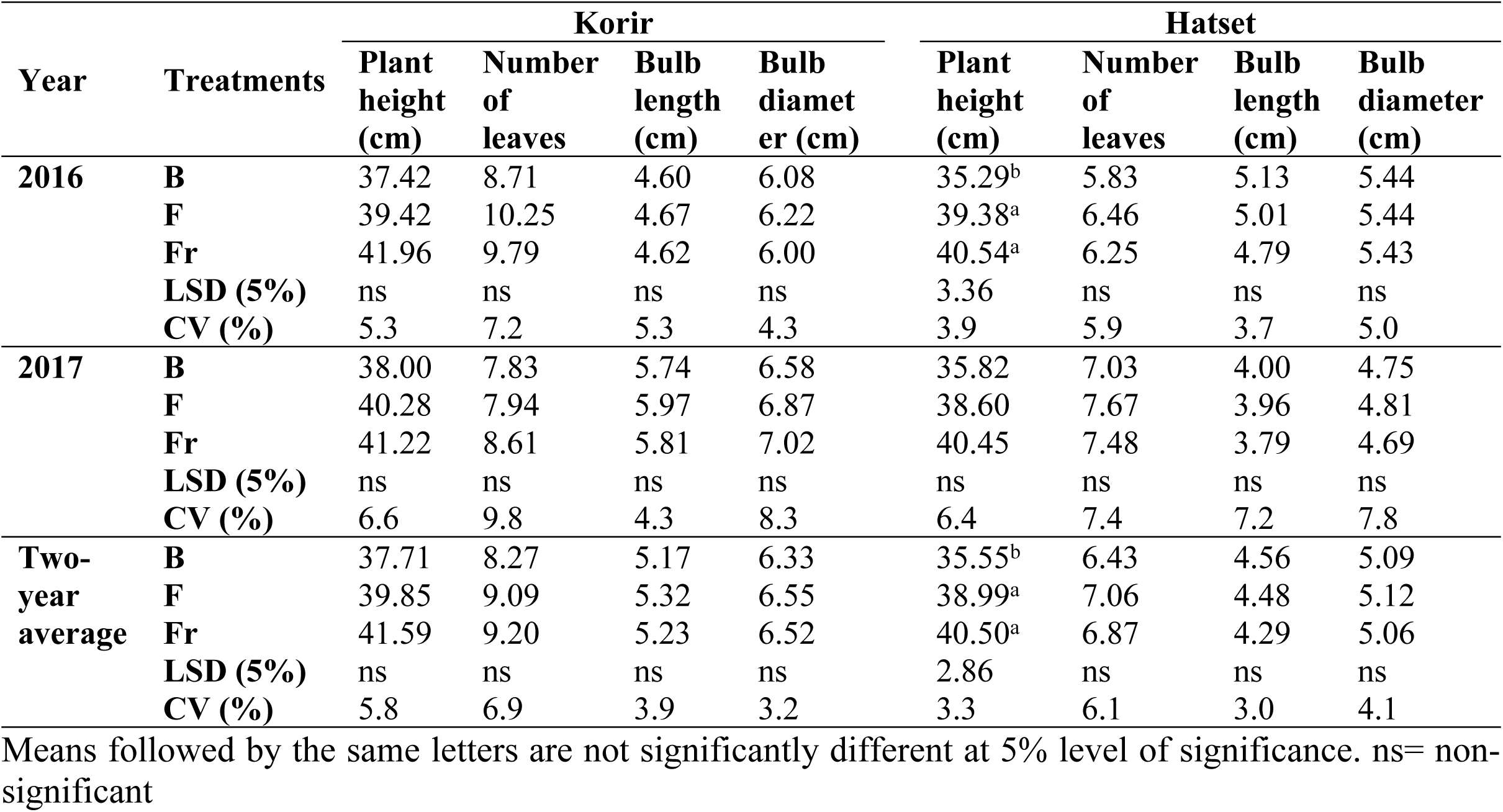
Effects of irrigation method on onion yield components.

**Table 9.**
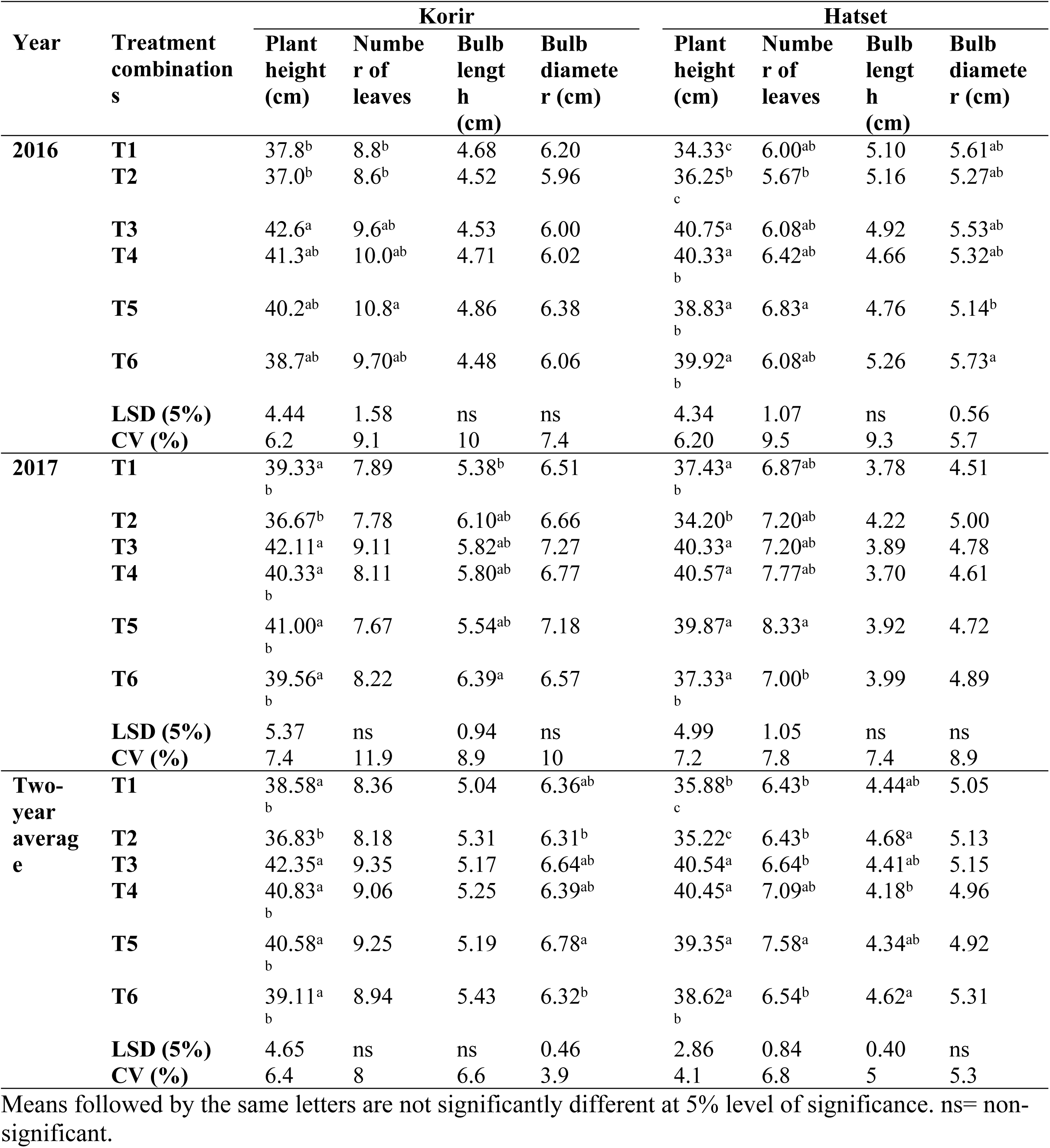
Factorial treatment combination effects of irrigation methods and irrigation interval on yield components at both sites.

| Year | Treatment combinations | Korir |  |  |  | Hatset |  |  |  |
| --- | --- | --- | --- | --- | --- | --- | --- | --- | --- |
|  |  | Plant height (cm) | Number of leaves | Bulb length (cm) | Bulb diameter (cm) | Plant height (cm) | Number of leaves | Bulb length (cm) | Bulb diameter (cm) |
| 2016 | T1 | 37.8<sup>b</sup> | 8.8<sup>b</sup> | 4.68 | 6.20 | 34.33<sup>c</sup> | 6.00<sup>ab</sup> | 5.10 | 5.61<sup>ab</sup> |
|  | T2 | 37.0<sup>b</sup> | 8.6<sup>b</sup> | 4.52 | 5.96 | 36.25<sup>b</sup><sub>c</sub> | 5.67<sup>b</sup> | 5.16 | 5.27<sup>ab</sup> |
|  | T3 | 42.6<sup>a</sup> | 9.6<sup>ab</sup> | 4.53 | 6.00 | 40.75<sup>a</sup> | 6.08<sup>ab</sup> | 4.92 | 5.53<sup>ab</sup> |
|  | T4 | 41.3<sup>ab</sup> | 10.0<sup>ab</sup> | 4.71 | 6.02 | 40.33<sup>a</sup><sub>b</sub> | 6.42<sup>ab</sup> | 4.66 | 5.32<sup>ab</sup> |
|  | T5 | 40.2<sup>ab</sup> | 10.8<sup>a</sup> | 4.86 | 6.38 | 38.83<sup>a</sup><sub>b</sub> | 6.83<sup>a</sup> | 4.76 | 5.14<sup>b</sup> |
|  | T6 | 38.7<sup>ab</sup> | 9.70<sup>ab</sup> | 4.48 | 6.06 | 39.92<sup>a</sup><sub>b</sub> | 6.08<sup>ab</sup> | 5.26 | 5.73<sup>a</sup> |
|  | LSD (5%) | 4.44 | 1.58 | ns | ns | 4.34 | 1.07 | ns | 0.56 |
|  | CV (%) | 6.2 | 9.1 | 10 | 7.4 | 6.20 | 9.5 | 9.3 | 5.7 |
| 2017 | T1 | 39.33<sup>a</sup><sub>b</sub> | 7.89 | 5.38<sup>b</sup> | 6.51 | 37.43<sup>a</sup><sub>b</sub> | 6.87<sup>ab</sup> | 3.78 | 4.51 |
|  | T2 | 36.67<sup>b</sup> | 7.78 | 6.10<sup>ab</sup> | 6.66 | 34.20<sup>b</sup> | 7.20<sup>ab</sup> | 4.22 | 5.00 |
|  | T3 | 42.11<sup>a</sup> | 9.11 | 5.82<sup>ab</sup> | 7.27 | 40.33<sup>a</sup> | 7.20<sup>ab</sup> | 3.89 | 4.78 |
|  | T4 | 40.33<sup>a</sup><sub>b</sub> | 8.11 | 5.80<sup>ab</sup> | 6.77 | 40.57<sup>a</sup> | 7.77<sup>ab</sup> | 3.70 | 4.61 |
|  | T5 | 41.00<sup>a</sup><sub>b</sub> | 7.67 | 5.54<sup>ab</sup> | 7.18 | 39.87<sup>a</sup> | 8.33<sup>a</sup> | 3.92 | 4.72 |
|  | T6 | 39.56<sup>a</sup><sub>b</sub> | 8.22 | 6.39<sup>a</sup> | 6.57 | 37.33<sup>a</sup><sub>b</sub> | 7.00<sup>b</sup> | 3.99 | 4.89 |
|  | LSD (5%) | 5.37 | ns | 0.94 | ns | 4.99 | 1.05 | ns | ns |
|  | CV (%) | 7.4 | 11.9 | 8.9 | 10 | 7.2 | 7.8 | 7.4 | 8.9 |
| Two-year average | T1 | 38.58<sup>a</sup><sub>b</sub> | 8.36 | 5.04 | 6.36<sup>ab</sup> | 35.88<sup>b</sup><sub>c</sub> | 6.43<sup>b</sup> | 4.44<sup>ab</sup> | 5.05 |
|  | T2 | 36.83<sup>b</sup> | 8.18 | 5.31 | 6.31<sup>b</sup> | 35.22<sup>c</sup> | 6.43<sup>b</sup> | 4.68<sup>a</sup> | 5.13 |
|  | T3 | 42.35<sup>a</sup> | 9.35 | 5.17 | 6.64<sup>ab</sup> | 40.54<sup>a</sup> | 6.64<sup>b</sup> | 4.41<sup>ab</sup> | 5.15 |
|  | T4 | 40.83<sup>a</sup><sub>b</sub> | 9.06 | 5.25 | 6.39<sup>ab</sup> | 40.45<sup>a</sup> | 7.09<sup>ab</sup> | 4.18<sup>b</sup> | 4.96 |
|  | T5 | 40.58<sup>a</sup><sub>b</sub> | 9.25 | 5.19 | 6.78<sup>a</sup> | 39.35<sup>a</sup> | 7.58<sup>a</sup> | 4.34<sup>ab</sup> | 4.92 |
|  | T6 | 39.11<sup>a</sup><sub>b</sub> | 8.94 | 5.43 | 6.32<sup>b</sup> | 38.62<sup>a</sup><sub>b</sub> | 6.54<sup>b</sup> | 4.62<sup>a</sup> | 5.31 |
|  | LSD (5%) | 4.65 | ns | ns | 0.46 | 2.86 | 0.84 | 0.40 | ns |
|  | CV (%) | 6.4 | 8 | 6.6 | 3.9 | 4.1 | 6.8 | 5 | 5.3 |

Means followed by the same letters are not significantly different at 5% level of significance. ns= non-significant.

#### Effects irrigation intervals on yield components of onion

As shown in Fig 4, a comparison of the irrigation intervals (fixed interval and farmers practices) was carried out for each of the onion growth parameters considered. As it is explicitly demonstrated by the strongly overlapped standard deviation error bars, the yield components including plant height, the number of leaves per plant, bulb length and bulb diameter was not significantly affected by the irrigation intervals for both sites and years (results not shown here). However, differences in parameters were clearly shown at both irrigation seasons and sites between the irrigation intervals. At the Korir site, the farmer’s practice (Fp) has shown better performance than the fixed irrigation interval (Fi) for the plant height, number of leaves and bulb diameter at both years and their mean. In contrast, considering the two-year average, the fixed irrigation interval has performed well in bulb length. Similarly, the performance of the fixed and farmers’ practice irrigation interval of the Hatset experimental site was also evaluated at each season (Fig 4). Accordingly, differences in the yield components of the onion crop were observed at both seasons due to irrigation scheduling differences. The irrigation water application, by farmers, revealed higher performances on the crop growth parameters such as plant height (except in 2016), the number of leaves per plant and bulb diameters than the fixed watering interval during each season and their mean. However, the fixed interval has shown relatively higher performance than the farmer’s practices in bulb length (Fig 4).

In summary, although statistically non-significant, the farmer’s practice showed better performance for all growth parameters except bulb length at both sites and irrigation seasons. This suggests that the use of a fixed irrigation interval is encouraging to save irrigation water. This indicates that farmers in the region may have better understandings of their irrigation lands. In summary, it is noticed that the growth parameters have shown better performance in both experimental sites. Similar results are also found in different studies that supports the argument [49,50].

#### Effects different irrigation methods on yield components of onion

The effects of irrigation methods on the majority of onion yield components have shown non-significant differences in both years and sites (Table 8). However, the two-year average result indicates that the furrow irrigation method (Fr) has shown higher plant height and number of leaves per plant. In terms of bulb length and bulb diameter, the flood irrigation method showed a higher average of the two-year values in both sites. In general, the plant height and number of leaves per plant showed higher results under the flood and furrow irrigation methods. However, higher bulb diameter and bulb length were obtained under flood and basin irrigation methods. Since the interest is to conserve water while keeping optimum onion productions, the basin and furrow irrigation methods are suggested for use by the farmers. These irrigation methods use techniques where water is conserved and used further to produce onion and other vegetable crops.

#### Treatment interaction effects on yield components of onion

Similar to the yield, the interaction effect of irrigation interval and irrigation methods were evaluated for the yield components. The analysis of the factorial interaction effects of irrigation methods and irrigation intervals on the yield components at yearly basis and an average of the two-year results are summarized in Table 9. The factorial interaction effects of the treatments showed a significant effect on plant height and number of leaves per plant. The highest plant height was obtained from treatment combination T3 and the least from T2 for each year and site. The highest leaf number was recorded from a treatment combination of T5 (10.8 at Korir, 2016) and the least was from T2 (5.67 at Hatset, 2016) for both sites (Table 9). For the bulb length, a significant interaction effect among the treatments is also found at Korir during 2017 season. As can be seen in Table 9, the higher bulb length is found in the factorial treatment combinations of T6 with 6.39 and 5.26 cm at Korir and Hatset sites, respectively.

When the yield components obtained from both growing seasons combined/averaged, a significant difference among the treatments were found for the plant height and bulb diameter at Korir site (Table 9). Similarly, except for the bulb diameter, all yield components have shown significant differences among the treatments in Hatset irrigation site. The observed variations between the experimental sites may be attributed to the soil characteristic variations observed in the irrigation schemes (Table 2).

### Effects of irrigation intervals and interactions with irrigation methods on water use and water productivity

#### Effects of irrigation intervals on water use and water productivity

Appropriate irrigation management practices are required to satisfy both high yield and higher water productivity. The irrigation scheduling techniques explained by fixed irrigation (Fi) and farmers’ practice (Fp) irrigation intervals are evaluated for their advantage of water productivity. The irrigation interval did not show a significant effect on total water productivity (WP) and irrigation water productivity (IWP) in both case studies. Fig 5 provides comparisons of irrigation intervals (based on crop water requirement and local practice) for the independent and the average of the irrigation seasons. The average result showed that applying a fixed irrigation interval exhibits a non-significant difference in the WP and IWP. In Korir irrigation site, the WP and IWP was 5.17 kg/m^3^ and 3.62 kg/m^3^ and 5 kg/m^3^ and 2.73 kg/m^3^ for Fi and Fp, respectively. Similarly, the WP and IWP at Hatset was 5.08 kg/m^3^ and 3.54 kg/m^3^ and 5.04 kg/m^3^ and 2.48 kg/m^3^ for Fi and Fp, respectively (Fig 5).

**Fig 5.**
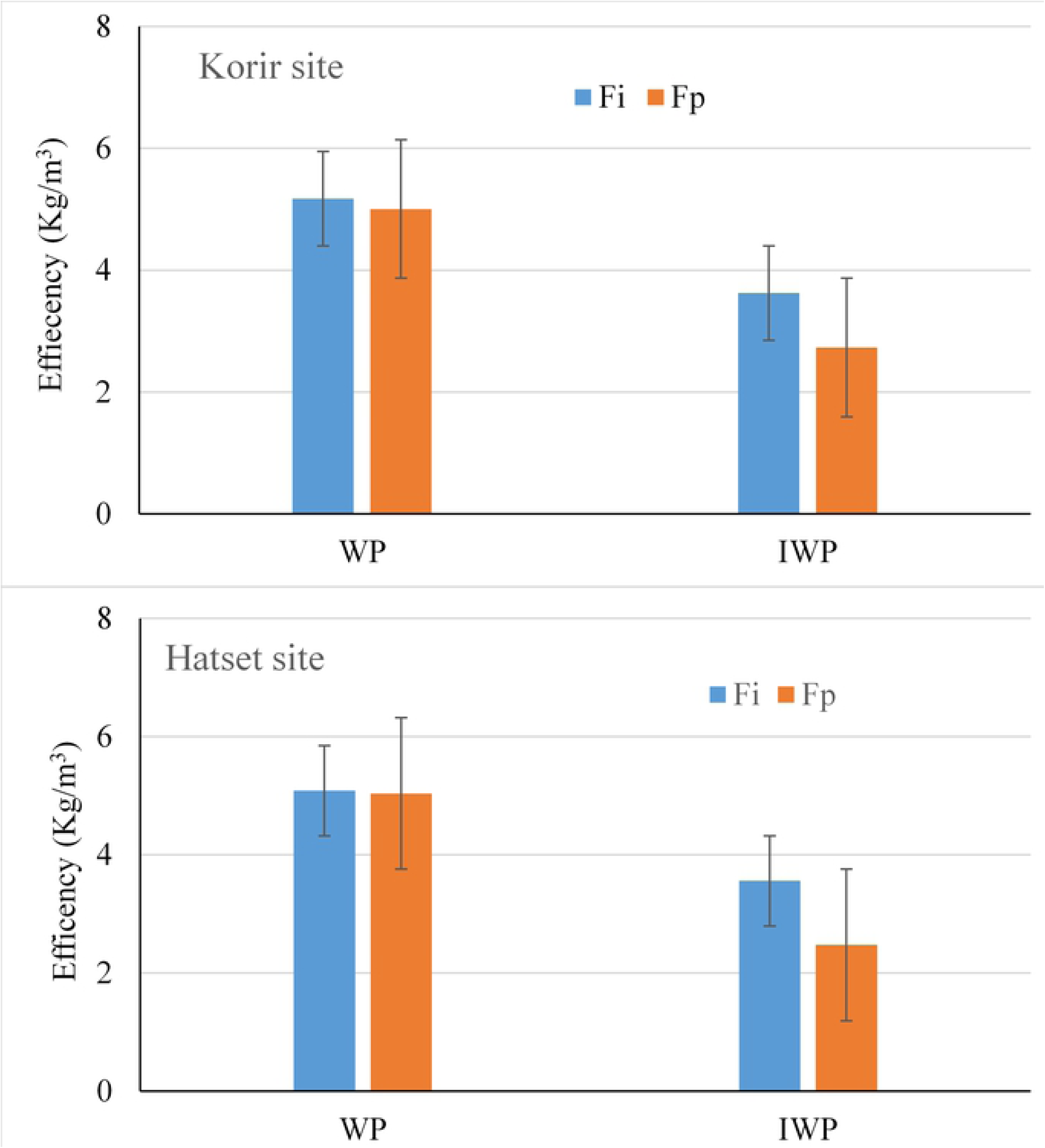
Effects of irrigation interval on onion WP and IWP in 2016.

#### Treatment interaction effects on water productivity

Table 10 shows the WP and IWP variations among the treatments during the 2016 and 2017 irrigation season at both sites. At the Korir experimental site, the highest and lowest WP were obtained from the factorial treatments of T1 (6.25 kg/m^3^) and T6 (3.72 kg/m^3^), respectively, during the 2016 growing season. In contrast, during the 2017 irrigation season, the highest and least WP was obtained from T2 and T5 treatments, respectively. A similar result was also observed in the Hatset irrigation scheme. The IWP was also significantly affected by the interaction effect of irrigation method and irrigation interval. For both years and sites, the highest and lowest IWP was obtained from treatment T2 (4.63 kg/m^3^) and the least was from T5 (2.08kg/m^3^), respectively. Similarly, combining the two years result, statistically significant differences were observed among the treatment combinations in both sites (Table 10). The lower WP and IWP values were recorded in T3 treatment.

**Table 10.** Factorial treatment combination effects of irrigation methods and irrigation interval on IWP and WP of onion.

| Year | Treatment combinations | Korir |  | Hatset |  |
| --- | --- | --- | --- | --- | --- |
|  |  | WP (kg/m<sup>3</sup>) | IWP (kg/m<sup>3</sup>) | WP (kg/m<sup>3</sup>) | IWP (kg/m<sup>3</sup>) |
| 2016 | T1 | 6.25<sup>a</sup> | 2.94<sup>b</sup> | 6.10<sup>a</sup> | 2.99<sup>cd</sup> |
|  | T2 | 5.93<sup>a</sup> | 4.15<sup>a</sup> | 6.02<sup>ab</sup> | 4.22<sup>a</sup> |
|  | T3 | 4.34<sup>a</sup> | 2.41<sup>bc</sup> | 5.11<sup>bc</sup> | 2.51<sup>de</sup> |
|  | T4 | 4.17<sup>b</sup> | 2.92<sup>b</sup> | 4.91<sup>c</sup> | 3.44<sup>bc</sup> |
|  | T5 | 4.37<sup>b</sup> | 2.12<sup>c</sup> | 4.79<sup>c</sup> | 2.36<sup>e</sup> |
|  | T6 | 3.72<sup>b</sup> | 2.60<sup>bc</sup> | 5.03<sup>c</sup> | 3.52<sup>b</sup> |
|  | LSD (5%) | 1.50 | 0.65 | 0.93 | 0.51 |
|  | CV (%) | 1.72 | 1.25 | 0.96 | 0.88 |
| 2017 | T1 | 5.98<sup>ab</sup> | 3.26<sup>bc</sup> | 6.30<sup>a</sup> | 3.10<sup>b</sup> |
|  | T2 | 6.61<sup>a</sup> | 4.63<sup>a</sup> | 6.10<sup>a</sup> | 4.27<sup>a</sup> |
|  | T3 | 4.34<sup>b</sup> | 2.37<sup>c</sup> | 3.68<sup>b</sup> | 1.81<sup>d</sup> |
|  | T4 | 4.67<sup>b</sup> | 3.27<sup>bc</sup> | 3.92<sup>b</sup> | 2.75<sup>bc</sup> |
|  | T5 | 4.74<sup>b</sup> | 2.59<sup>c</sup> | 4.24<sup>b</sup> | 2.08<sup>cd</sup> |
|  | T6 | 5.93<sup>ab</sup> | 4.15<sup>ab</sup> | 4.52<sup>b</sup> | 3.16<sup>b</sup> |
|  | LSD (5%) | 1.83 | 1.17 | 1.41 | 0.84 |
|  | CV (%) | 1.87 | 1.91 | 1.62 | 1.75 |
| Two-year average | T1 | 6.11<sup>a</sup> | 3.33<sup>b</sup> | 6.20<sup>a</sup> | 3.05<sup>b</sup> |
|  | T2 | 6.27<sup>a</sup> | 4.39<sup>a</sup> | 6.06<sup>a</sup> | 4.24<sup>a</sup> |
|  | T3 | 4.34<sup>b</sup> | 2.37<sup>d</sup> | 4.39<sup>b</sup> | 2.16<sup>c</sup> |
|  | T4 | 4.42<sup>b</sup> | 3.10<sup>bc</sup> | 4.42<sup>b</sup> | 3.09<sup>b</sup> |
|  | T5 | 4.56<sup>b</sup> | 2.48<sup>cd</sup> | 4.52<sup>b</sup> | 2.22<sup>c</sup> |
|  | T6 | 4.82<sup>b</sup> | 3.37<sup>b</sup> | 4.77<sup>b</sup> | 3.34<sup>b</sup> |
|  | LSD (5%) | 1.13 | 0.72 | 0.94 | 0.57 |
|  | CV (%) | 1.22 | 1.25 | 1.02 | 1.04 |

Means followed by the same letters are not significantly different at 5% level of significance. ns= non-significant.

## Discussion

The effects of irrigation intervals on onion yields produced higher yield under Fi is as compared to the Fp, though non-significant. Generally, both irrigation intervals (Fi and Fp) have shown higher yield productions. In terms of the onion yields obtained from both intervals, different studies have reported similar results elsewhere in the world [51,52]. For instance, the application of six days irrigation interval has resulted in 136.6 q/ha bulb yield [51]. Gwandu and Idris [52] showed that irrigating every seven days on sandy loam soils can give 190.8 kg/ha of bulb yield. Bossie et al. [22] indicated that the use of a fixed irrigation interval instead of the customary water delivery system could further encourage farmers. Moreover, farmer irrigation practices have been reported to increase water usage for a given crop yield [16,45,50]. A fixed irrigation interval could also be beneficial to use more crop per drop of water enhancing the greater value of water in terms of water prices [14]. In summary, the fixed irrigation interval has a multitude of benefits to onion producing farmers as it contributes better water management, reduces waterlogging and salinity in the schemes [16,18]. Majority of onion yield components were not significantly affected by the irrigation intervals in both sites. However, some of the parameters (e.g., plant height and the number of leaves per plant) were significantly affected during the interaction effect in Hatset site. The observed variations between the experimental sites may be attributed to the soil characteristic variations observed in the irrigation schemes (Table 2).

For the effects of irrigation methods on onion yield, the use of basin irrigation method produces a higher yield of onion crop. Similar results also reported that the basin irrigation method produces higher onion yields as compared to the other irrigation methods [25,45,53,54]. Moreover, the result is also in line with the farmers’ irrigation methods preferences. Farmers in most irrigated areas of Tigray usually apply basin irrigation method to their onion crop cultivation. It is noteworthy to explain that the extension experts in the region recommend to use furrow irrigation for onion production which contradicts this result and farmers preferences. It is suggested that improved irrigation methods have better benefits in agricultural water management works in semiarid irrigation schemes [30].

Furthermore, this study demonstrated that improved irrigation scheduling techniques can improve water productivity. This suggests that for increased WP and IWP the improved irrigation scheduling technologies could have paramount importance. Inline to our results, different authors including, Belay et al. [25] and Mintesinot et al. [17] found similar results elsewhere in Ethiopia. Hence, conserving water resources are necessary to assure the economic and environmental sustainability of irrigated agriculture by consuming less water while maximizing yield. These eventually improve irrigation scheduling techniques to produce higher onion yields and water productivity. Irrigation scheduling technologies that conserve water are also necessary to assure the economic and environmental sustainability of irrigated agriculture [55]. Effective water-saving irrigation strategies can improve both the crop yields and water productivity [56].

The key finding of this study is that a basin irrigation method with a fixed irrigation interval performed well both in terms of market interest and mass production of onion. The interaction results confirm that it is possible to increase onion productivity by applying these strategies. The results of onion production in the experimental sites are in agreement with some studies around the world [17,25,54,57]. These literatures reported a consistent result that the basin irrigation method gave a higher onion yield and optimum size of the onion bulb. Onion production in the region is delivered to the markets and the buyers in urban areas prefer a firm medium-sized onion bulb. Basin irrigation with fixed irrigation interval strategies was found to be a useful practice for better scheduling of onion with optimum production. Our result is more than 263q/ha from both sites which are even higher than that of reported in other studies [16,54]. However, the total onion production from the three irrigation methods is lower as compared to drip irrigation and sprinkler irrigation systems [45]. This is due to the application of irrigation water using drip or sprinkler is superior in terms of water-saving and overall in a benefit-cost ratio of production [16,58]. Drip irrigation can double the yield and reduce the amount of water by halve but produce larger onion size compared to the furrow, basin and flood irrigation methods. This is also supported by Enciso et al. [55] who reported the size of onion irrigated by drip irrigation was greater than the size of furrow irrigated onion by more than 181%. On the other hand, irrigation methods such as furrow, flood and furrow are common in regions with layouts of small fields and the northern Ethiopia region is typically characterized by small irrigation schemes where farmers’ landholding capacity is not more than 0.25 ha [14,16,18,25].

In summary, farmers in the region are encouraged to use a basin and fixed interval irrigation methods to produce good marketable onion bulbs. Given the size of the plot to be irrigated is very small, the regional government also encourages farmers to grow high-value marketable cash crops such as onions. However, cautions need to follow when applying water in the basin irrigation as farmers may apply excess water. The results of this study could be considered as a guiding principle in deciding which irrigation method and interval should be used if the onion crop has to be highly productive under existing conditions in the Tigray region.

## Conclusions

A field experiment was carried out to evaluate irrigation methods and irrigation scheduling on onion productivity in the semiarid region of Ethiopia. The experiment was conducted in two irrigation sites for two consecutive years (2016 and 2017). This study was conducted in an experimental treatment arranged in a randomized complete block design at which plots were replicated three times. The results showed that the variations between the fixed irrigation intervals are non-significant in both sites and seasons. In contrast, irrigation methods showed significant differences in all conditions. The basin irrigation method and its corresponding factors from the irrigation interval treatments obtained higher performances with a total bulb yield of 263.85 q/ha and an irrigation water productivity of 4.32 kg/m^3^. Basin irrigation methods with fixed irrigation interval produced higher bulb yield in small irrigation schemes. Furthermore, the basin method showed that good marketable onion bulbs with firm medium-size as compared to the flood and furrow irrigation methods. This is important as the consumers in urban areas are more interested in medium-size onion bulb for their daily use. It can be concluded that improved WP and IWP can be achieved by applying basin irrigation with fixed irrigation interval under existing farmers’ landholding in the semiarid areas in northern Ethiopia. In addition, improved water management techniques can enhance water use efficiency and crop yield in areas with limited water resources. Results can be used as a guiding principle in deciding which irrigation method and interval can be used, particularly for a higher quantity and quality of onion production in areas where farmers have a very limited plot of irrigated land. This could enhance farmers’ income beyond their daily food security enhancements under the existing markets and onion productivity conditions in Tigray.

## Acknowledgements

The authors acknowledge the Tigray Agricultural Research Institute for financing the research work. We are also thankful to the administration and logistics of the Mekelle Agricultural Research Centre for their greater contribution towards the timely accomplishment of the research. Besides, the authors would like to thank for the Mekelle Soil Research Centre, laboratory technicians.

